# Cortical involvement in stroke survivors for balance maintenance

**DOI:** 10.64898/2026.02.18.706534

**Authors:** Thomas Legrand, Giorgia Pregnolato, Scott Mongold, Pierre Cabaraux, Antonella Ianotta, Andrea Baroni, Annibale Antonioni, Marc Vander Ghinst, Mathieu Bourguignon, Sofia Straudi, Giacomo Severini

## Abstract

Stroke survivors often experience balance impairments which may result in increased risk of falling. However, it is unclear whether and how a stroke changes cortical involvement for postural control. To clarify this issue, we assessed the effect of stroke on sway-based corticokinematic coherence (CKC), which is a measure of the coupling between cortical electrophysiological signals and postural sways. To that end, we recorded the center-of-pressure fluctuations and electroencephalographic cortical activity of 34 stroke survivors and 34 healthy participants performing balance tasks during which sensory information was manipulated, by either removal or alteration. We found significantly increased CKC derived from medio-lateral sway in stroke participants when standing on foam compared to healthy controls, suggesting an increase in cortical involvement to compensate for the decreased function of the hemiparetic side, even in highly functional stroke survivors. Moreover, a relationship was found between CKC and clinical scores (Berg Balance Scale and Fugl-Meyer). This suggests that CKC could be used as a biomarker to track progress beyond traditional functional recovery as measured by clinical scores.

## Introduction

Stroke, resulting from an interruption of cerebral blood flow due to arterial occlusion or hemorrhage, remains a leading cause of long-term disability worldwide, despite progress in acute management and preventive care (1, 2). One of the functional consequences following stroke is impairment in postural control, which compromises autonomy and increases the risk of falls in stroke survivors (3–5). For instance, the weight distribution on the lower limbs is asymmetric, with a shift favoring the unaffected limb (6, 7), affecting weight transfer and postural adjustments (8–10). As such, even stroke survivors with good functional recovery of the lower limbs may exhibit persistent impaired postural control (11). This suggests that maintaining postural control in daily life is not merely a matter of motor recovery but also relies on the availability of cortical resources and the efficient engagement of distributed brain networks. It is therefore critical to clarify the central mechanisms underlying the reorganization of postural control in this population, so that rehabilitative efforts align with the neural processes that support adaptive sensorimotor functioning and postural stability after stroke.

Already, previous work in stroke has highlighted several structural and functional alterations in the affected hemisphere, including: altered cortical thickness (12), changes in excitation-inhibition balance (13), and a reconfiguration of brain networks dependent, in part, by the extent of anatomical damage (14). In addition to these key alterations, diverse mechanisms that contribute to balance dysfunction in stroke have been identified (15), including motor weakness, asymmetrical muscle tone, sensory loss, and altered cognition. Yet, neurophysiological processes related to postural control remain underexplored. On the other hand, in the context of upper limb motor recovery, which dominates the stroke literature, impaired cortical sensorimotor processing and integration consistently emerge as a hallmark of stroke, even in the chronic phase (16). Indeed, there is evidence to suggest that this impaired sensorimotor function underlies poor postural control in stroke, at least from behavioral experiments. Dual-task paradigms that combine cognitive and postural challenges reveal that stroke survivors experience disproportionate decrements in stability, reflecting limited capacity to flexibly allocate cortical resources (12, 14). It can be argued that most postural adjustments are generated reflexively at the level of the spinal cord and brainstem; however, recent reports indicate that the cerebral cortex intervenes intermittently to adapt the postural corrective strategy to the environment (17, 18). This suggests that the amount of cortical resources available after a stroke may not be enough to properly maintain balance in all situations of daily living. Moreover, somatosensory deficits are prevalent in stroke survivors, affecting up to 89% of post-stroke individuals (19, 20) and are associated with motor deficits and reduced functional mobility (21, 22). Because maintaining upright posture requires the continuous integration of sensory information from visual, vestibular, and somatosensory systems (23, 24), there is a clear need to better understand the interplay between limited cortical resources and impaired sensory integration underlying postural control after stroke.

To assess sensorimotor integration for postural control, experimental paradigms have employed sensory manipulations, such as visual occlusion, while the participants perform balance tasks (25). These perturbations allow the assessment of the central nervous system’s responsiveness to altered sensory input. Combined with electroencephalography (EEG), which allows recording of cortical activity during weight-bearing tasks, studies have highlighted cortical activity time-locked to naturally occurring postural sway (18, 26, 27). Similarly, significant coherence, an extension of Pearson’s correlation to the frequency domain, between cortical activity and postural sway was found especially salient when sensory inputs were altered (28). More importantly, this coupling, termed corticokinematic coherence (CKC), was observed to be behaviourally relevant and was correlated with the modulation in postural sway brought about by the sensory manipulation of the balance task (28). These results suggest that CKC could be used to directly assess the magnitude of cortical involvement in postural control. Indeed, growing evidence indicates that, given the complexity of post-stroke reorganization mechanisms, combining multiple sources of information to capture both central and peripheral events represents an effective strategy for capturing the multidimensional nature of motor recovery (29, 30). Such an approach has the potential to improve the characterization of functional impairments but also enhances the ability to predict recovery trajectories, which is of particular relevance in this population.

In this study, we investigated the cortical involvement in balance control among stroke survivors and healthy controls by examining CKC during quiet stance under varying sensory conditions and along both the antero-posterior and medio-lateral axes. We hypothesize that sway-based CKC can capture differences in cortical involvement post-stroke, and that these differences relate to behavioral measures of balance performance and clinical scores. Understanding these mechanisms may help identify neurophysiological markers of functional recovery and guide the development of targeted rehabilitation strategies.

## Material and Methods

### Participants

The pool of participants is composed of 3 cohorts recruited separately, who performed the same balance protocol. The first cohort (*n_stroke_* = 8; *n_control_* = 8) was recruited at University College Dublin (UCD), the second (*n_stroke_* = 14) at Ferrara Rehabilitation University Hospital, and the third at Erasme University Hospital (*n_stroke_* = 12; *n_control_* = 26). The study had prior approval by the ethics committees of each recruiting centre (UCD: LS-23-64-Legrand-Sev, Dublin, Ireland; Ferrara: 190/2024/Oss/AOUFe, Area Vasta Emilia Centro Ethics Committee, Ferrara, Italy; Erasme: B4062021000323, Brussels, Belgium). Volunteers unable to stand or walk without the aid of an assistive device or suffering from co-morbidities affecting balance or locomotion were excluded from the study.

### Procedure

All stroke participants were assessed using the Berg Balance Scale (31) to evaluate their static and dynamic balance function. The Berg Balance Scale is a validated scale consisting of 14 items, with a score between 0 (no balance function) up to 56 (complete balance function). Participants recruited at UCD and Ferrara (*n* = 22) were also assessed using the Fugl-Meyer Assessment (FMA) (32), lower limb motor and sensation section, to collect information about their sensorimotor function. The FMA score is a 3-point clinical scale (0 = no function; 1 = partial function; 2 = normal function) with a maximum score for the lower limb of 34 points; sensation, with a maximum score of 24 points; and passive joint motion and pain, with a maximum score of 48 points. Both the Berg Balance Scale and FMA were administered by trained physiotherapists.

All participants, both stroke survivors and controls, performed 4 standing conditions, which included: bipedal quiet standing on a firm surface with eyes open (i) or closed (ii), or on a foam pad with eyes open (iii) or closed (iv). Each condition was recorded and lasted 5 minutes. Participants performed all conditions barefoot, without speaking, with their arms extended alongside their bodies, looking straight ahead at a cross on the wall facing them. The position of the feet was standardized, parallel at shoulder width, for all conditions. Participants were supervised by an assessor positioned immediately behind them to prevent falls to the ground. No contact was made between the participants and the assessor unless a fall would have occurred without assistance. No assistance was required across cohorts. Participants were monitored for self-reported fatigue and took mandatory breaks between trials.

### Data acquisition

Cortical activity was recorded during the balance tasks with a 128-channel EEG headcap (BioSemi, ActiveTwo, Amsterdam, Netherlands) at UCD, a 32-channel EEG headcap (Easy Cap GmbH, Herrsching, Germany) at Ferrara Rehabilitation Hospital, and a 63-channel EEG headcap (Waveguard original, ANT Neuro, Hengelo, Netherlands) at Erasme University Hospital. Electrodes, embedded in a cap, were arranged according to the 10/20 system. Impedances of the electrodes were kept below 50 kΩ using electrolyte gel, and the reference was set at CPz. EEG signals were sampled at 1000 Hz. Additionally, ground reaction forces and moments were recorded at 1000 Hz during each condition with a force plate (UCD: Model 4060, BERTEC, Columbus, OH, USA; Ferrara: Model 4080-10, BERTEC, Columbus, OH, USA; Erasme: AccuSway-O, AMTI, Watertown, MA, USA).

### Data pre-processing

EEG and force plate data were imported into Matlab (Mathworks, Natick, MA, USA). After examination of the raw data in all participants, EEG electrodes on the mastoids (M1 and M2), when present, were removed because they featured high-amplitude artifacts caused by poor or unstable skin-electrode contact. Thus, signals from the remaining electrodes were kept for further analysis.

EEG data were band-pass filtered between 0.2 and 90 Hz and notch filtered at 50 Hz using a fourth-order Butterworth filter. Bad channels were identified based on the recommendations provided by Bidgely-Shamlo et al. (33). Signals from bad electrodes were interpolated based on the signals of the surrounding electrodes (34). EEG signals were then re-referenced to their common average. Finally, 20 independent components were evaluated from the data with Fast ICA (35) to identify and suppress further physiological artifacts. Independent components corresponding to heartbeat, eye-blink, and eye movement artifacts were visually identified, and corresponding signals reconstructed by means of the mixing matrix were removed from the full-rank data.

Force plate data were band-pass filtered between 0.1 and 150 Hz with a fourth-order Butterworth filter.

## Data analysis

### Center of Pressure (CoP) computation

CoP time-series, antero-posterior (CoP_AP_), and medio-lateral (CoP_ML_) were filtered between 0.1 and 10 Hz. The Euclidean norm of CoP_AP_ and CoP_ML_ at every time step yielded the excursion (rCoP). The single derivative of the CoP time-series yielded the CoP velocities (vCoP_AP_ and vCoP_ML_).

We also estimated the standard deviation of the CoP along the medio-lateral (SD(CoP_ML_)) and antero-posterior axes (SD(CoP_AP_)) and the mean of the Euclidean norm of CoP velocity (mean(|vCoP|)) to quantify the amount of postural sway during a standing condition.

### CKC computation

CKC was assessed using coherence analysis. Coherence is an extension of the Pearson correlation coefficient to the frequency domain, which quantifies the degree of coupling between two signals, i.e., CKC strength, by providing a number between 0 (no linear dependency) and 1 (perfect linear dependency) for each frequency (36). In practice, EEG and CoP data were divided into overlapping 2-s epochs with 1.6-s epoch overlap (leading to a frequency resolution of 0.5 Hz). We retained only the 4 electrodes from among those typically overlying the lower limb sensorimotor area (SM1) and common to the 3 different EEG headcaps used for recording (Cz, C3, C4, Fz). To ensure comparability between conditions, some epochs were discarded to reach a similar number of artifact-free epochs in all four conditions within each participant (534 ± 6; mean ± SD across participants). The discarded epochs were the last ones of the last recording block. The retained epochs were then Fourier-transformed and combined to derive a spectrum of coherence between the CoP features (CoP_ML_, rCoP, and vCoP_AP_) and each retained EEG signal, following the formulation of Halliday et al. (36), and using the multitaper approach (3 orthogonal Slepian tapers, yielding a spectral smoothing of 1.5 Hz) to estimate power- and cross-spectral densities (37). The CoP features rCoP and vCoP_AP_ were selected because they were successfully detected in young healthy participants and seemed to relate to two different pathways, efferent and afferent, respectively (28). Additionally, CoP_ML_ was selected, as stroke survivors often display postural asymmetry (11) and could thus unveil differential cortical involvement related to asymmetrical weight distribution during postural control.

For further analyses, we retained only the electrode that yielded the highest CKC averaged over balance conditions and across 0.5-10 Hz for each feature. From this point on, it is referred to as EEG_SM1_. CKC strength was estimated as the mean CKC value across 0.5-10 Hz at the EEG_SM1_ electrode.

### Time delay estimation

Time delays were estimated from the phase slope of the cross-spectral density between the EEG_SM1_ signal and each CoP feature, using only the data of participants with significant coherence. The phase-slope for a given participant and CoP feature was estimated at frequencies where significant coherence was found within the range 0.5-10 Hz. A minimum of 6 consecutive significant frequency bins was required to compute a linear regression. A negative-phase slope indicated that the cortical activity was led by the CoP variation. Time delays were also estimated between CoP features using the same approach.

### Statistical analysis

Statistical analyses were performed using Matlab (Mathworks, Natick, MA, USA).

A threshold for statistical significance of CKC (*p* < 0.05 corrected for multiple comparisons) was obtained as the 95^th^ percentile of the distribution of the maximum coherence (across 0.5 – 10 Hz, and across the SM1 sensor selection) evaluated between EEG and Fourier transform surrogate reference signals (1000 repetitions) (38). The Fourier transform surrogate of a signal is obtained by computing its Fourier transform, replacing the phase of the Fourier coefficients by random numbers in the range [-π; π], and then computing the inverse Fourier transform (38).

Normality of the data was assessed using the Kolmogorov-Smirnov test. When the data was found not to be normally distributed, a Box-Cox transformation was applied prior to subsequent analyses.

A linear mixed-effect model was used to assess the effect of the balance condition and of a stroke on mean(|vCoP|), SD(CoP_ML_), SD(CoP_AP_), and CKC strength for each feature. The fixed effects were defined as vision (eyes open or closed), surface (firm or foam), and health status (control or stroke), and their interactions. In order to take into account the variability between the 3 cohorts, due to differences in recording equipment and participants’ demographics, the collection site (UCD, Ferrara, and Erasme) was added as a random effect in the model, allowing for random intercepts and slopes for the fixed effects. The effective degrees of freedom of the t-statistics on the random effects were computed with the Welch-Satterthwaite equation. Post-hoc tests were performed when significant fixed effects were found, and the alpha level was corrected with the Bonferroni criterion.

Multiple regression analyses were conducted between the average of standardized CoP postural-sway parameters (SD(CoPAP) and mean(|vCoP|)) and CKC strength for the 3 CoP features across all balance conditions, separately for the 2 groups. To ease the interpretation of regression weights, values of CKC strength were standardized.

We assessed the association between the Berg Balance Scale and CKC, as well as between clinical scores and postural sway parameters (SD(CoP_ML_), SD(CoP_AP_), and mean(|vCoP|)) for all standing conditions using linear effect models. The fixed effects were the balance conditions.

## Results

A cohort of 34 stroke participants (mean ± SD; age: 60 ± 15 y.o.; 12 women) and 34 healthy controls (age: 59 ± 18 y.o.; 18 women) were recruited. Among the 34 stroke participants, 28 were at a chronic stage (> 6 months after stroke) and 6 were in the sub-acute phase after stroke (< 6 months), 19 presented with hemiparesis on the left side and 12 on the right side. Additional details about the type of stroke and lesion locations are provided in supplementary information (Table S1).

### Clinical outcomes

The results of the clinical scales are presented in Table 1. The Berg Balance Scale scores indicated an overall good balance and functional mobility for our stroke cohort, with the exception of 1 participant with a score < 41. The FMA - lower extremity scores indicated, on average, a high level of mobility function (39), with only one participant scoring 20 (moderate impairment). Eight participants presented sensation impairment, and 7 had some limitations in the passive range of motion. Only one participant reported pain.

**Table 1.** Results of the clinical tests for the stroke participants.

| | score (mean ± SD) | range |
| --- | --- | --- |
| Berg Balance Scale ($n = 34$) | | |
| | $52.7 \pm 5$ | [32 - 56] |
| Fugl-Meyer Assessment - Lower extremity sensorimotor function ($n = 22$) | | |
| Lower extremity | 28.4 ± 4.2 | [20 - 34] |
| Sensation | 10.7 ± 2.4 | [3 - 12] |
| Passive joint motion | 19.1 ± 1.8 | [14 - 20] |
| Joint pain | 19.9 ± 0.9 | [17 - 20] |

### Postural sway

Figure 1 presents the distribution of the CoP standard deviation along the medio-lateral axis (SD(CoP_ML_)), antero-posterior axis (SD(CoP_AP_)) and the average CoP velocity (mean(|vCoP|)) for both control and stroke participants in each balance condition. A linear mixed-effect model applied to SD(CoP_ML_) unveiled a significant effect of vision (*F_1,265_* = 4.0, *p* = 0.047) and of having suffered a stroke (*F_1,265_* = 99.2, *p* < 0.001), along with a significant interaction between the two (*F_1,265_* = 4.4, *p* = 0.037). Applied to SD(CoP_AP_), the model unveiled a significant effect of vision (*F_1,265_* = 100.2, *p* < 0.001), surface (*F_1,265_* = 13.0, *p* < 0.001) and of having suffered a stroke (*F_1,265_* = 23.1, *p* < 0.001), along with significant double interactions (vision x surface: *F_1,265_*= 64.5, *p* < 0.001; vision x group: *F_1,265_* = 15.2, *p* < 0.001; surface x group: *F_1,265_* = 58.9, *p* < 0.001). Similar effects were found for mean(|vCoP|) for vision (*F_1,265_* = 40.0, *p* < 0.001) and surface (*F_1,265_*= 8.2, *p* = 0.004), but in this case having suffered a stroke did not have an impact (*F_1,265_* = 0.8, *p* = 0.362), while the interactions vision x surface (*F_1,265_* = 29.4, *p* < 0.001) and surface x group (*F_1,265_* = 24.1, *p* < 0.001) stayed significant. The random effects statistical outcomes are reported in Supplementary information (Table S1 and Table S2). Overall, the data collection site accounted for significant variations in both SD(CoP_AP_) and mean(|vCoP|). In particular, their grand averages were significantly different, and their relationship to vision or surface was significantly different (Table S2, S3, and S4). However, their relationship to having suffered a stroke was not significantly different. The results of post-hoc comparisons are presented in Figure 1A for SD(CoP_AP_) and Figure 1B for mean(|vCoP|). Overall, sways were larger when standing with eyes closed compared with eyes open, and when standing on foam blocks compared with a firm surface for both groups (Figure 2). Standing on foam with the eyes closed led to an even larger significant increase irrespective of health status (Figure 1). When standing on a firm surface, eyes closed, stroke participants displayed significantly higher mean(|vCoP|) compared to the control group (Figure 1B).

**Figure 1.**
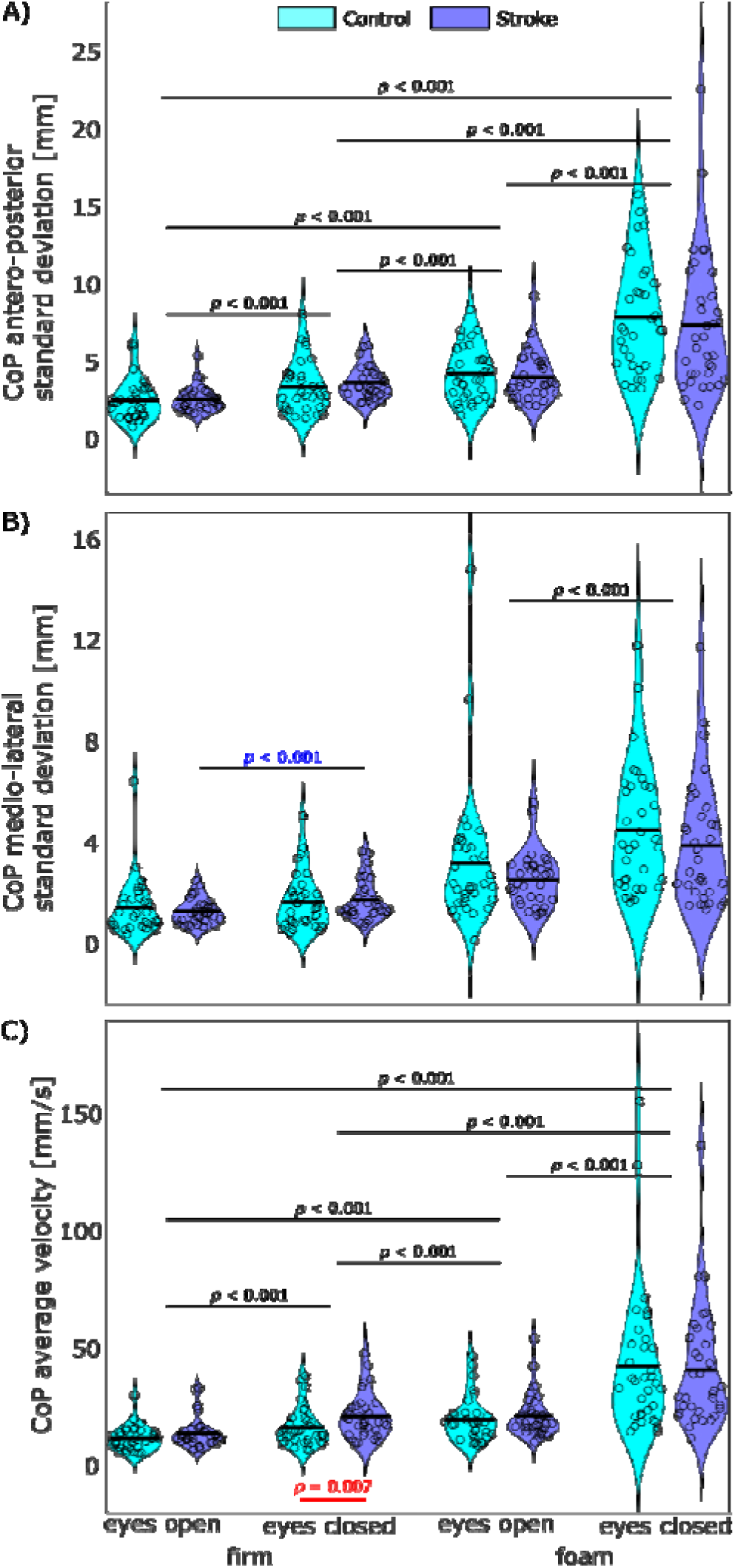
Distribution of instability parameters in the 4 conditions and 2 groups. A — Distribution of CoP_AP_ standard deviation. B — Distribution of CoP_ML_ standard deviation. C — Distribution of the average CoP velocity. Each individual participant’s value is indicated with circles and a horizontal black bar indicates the group average. Significant differences between groups (in red) and between conditions (in black when the results are identical for both groups and in color corresponding to the group otherwise) are indicated with horizontal lines.

**Figure 2.**
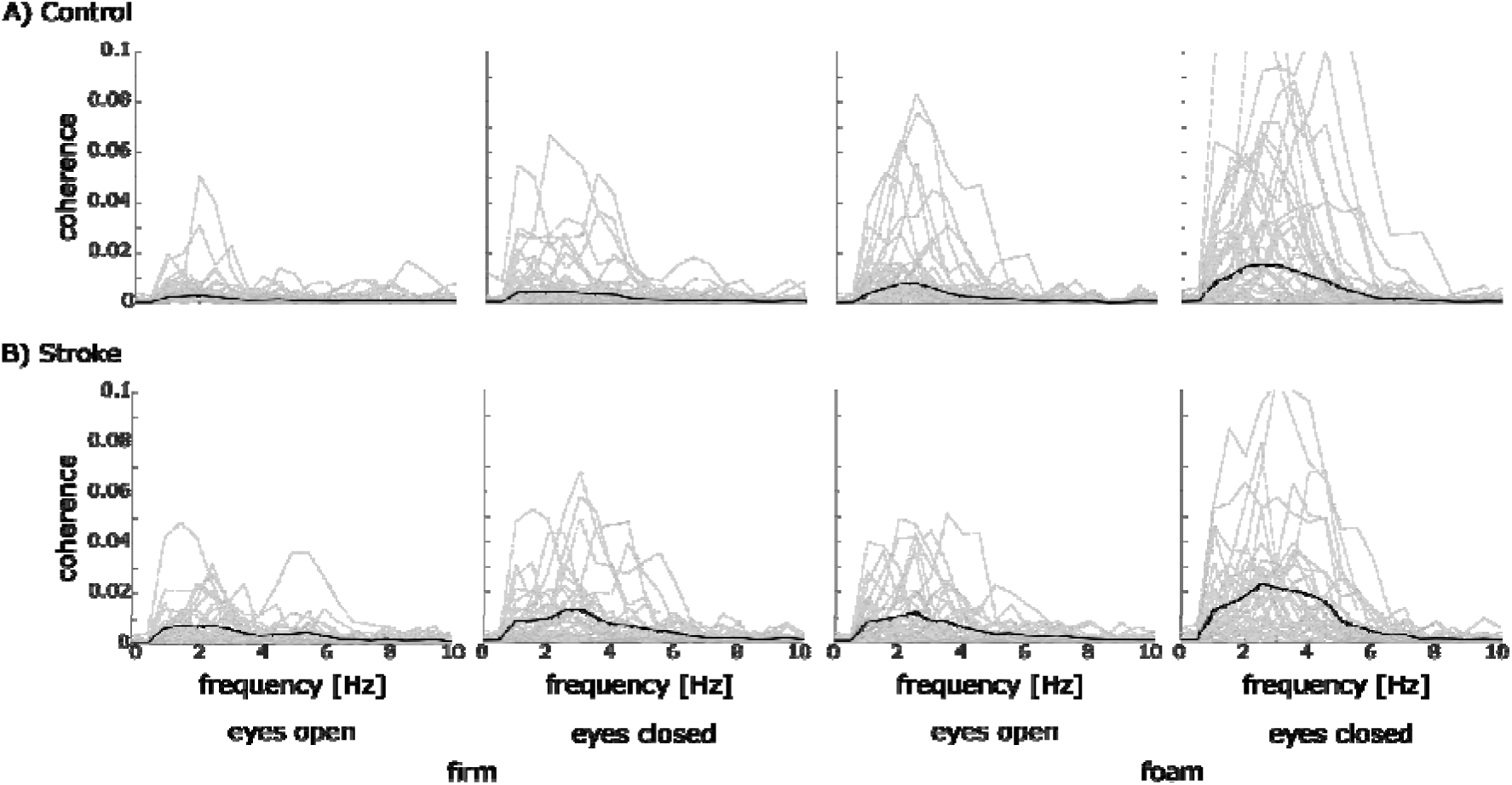
Sway-based CKC spectra derived from rCoP at the maximum electrode covering the primary sensorimotor area for each group and for all standing conditions. The trace of each individual is displayed in light gray while the group average is in black.

### Cortical encoding of CoP

Table 2 presents the number and percentage of participants displaying significant CKC, for each feature, age group and balance condition. Qualitatively, across all CoP features, the proportion of participants with significant CKC increased with condition complexity and was larger in stroke participants when standing on a firm surface with eyes open, whereas it was smaller when standing on foam, with or without eyes open.

**Table 2:** Number and percentage of participants with significant CKC with each feature.

| CKC | surface | firm |  | foam |  |
| --- | --- | --- | --- | --- | --- |
|  | eyes | open | closed | open | closed |
| $vCOP_{AP}$ | Control | 10 (29%) | 9 (26%) | 14 (41%) | 23 (68%) |
|  | Stroke | 11 (32%) | 12 (35%) | 12 (35%) | 17 (50%) |
| $COP_{ML}$ | Control | 2 (6%) | 2 (6%) | 3 (9%) | 4 (12%) |
|  | Stroke | 4 (12%) | 3 (9%) | 5 (15%) | 9 (26%) |
| rCOP | Control | 2 (6%) | 8 (24%) | 11 (32%) | 21 (62%) |
|  | Stroke | 3 (9%) | 6 (18%) | 8 (24%) | 17 (50%) |

Figure 2 presents the CKC spectra derived from |rCoP| for both groups in all standing conditions. Across participants, CKC tended to peak between 0.5 and 6 Hz.

Figure 3 presents the distribution of individual CKC strength (CKC values at the peak electrode around Cz, averaged across 0.5-10 Hz) for each CoP feature. The linear mixed effect model revealed a significant effect of the standing surface (*F_1,265_* = 6.1, *p* = 0.014) and of having suffered a stroke (*F_1,265_*= 1910.9, *p* < 0.001) with significant interactions (vision x surface: *F_1,265_* = 80.7, *p* < 0.001; vision x group: *F_1,265_* = 25.7, *p* < 0.001; surface x group: *F_1,265_* = 19.3, *p* < 0.001; vision x surface x group: *F_1,265_* = 111.4, *p* < 0.001) on CKC derived from rCoP. Concerning the CKC derived from CoP_ML_, vision (*F_1,265_* = 511.9, *p* < 0.001), surface (*F_1,265_*= 60.9, *p* < 0.001), having suffered a stroke (*F_1,265_*= 1724.9, *p* < 0.001) were significant, as well as all their interactions (vision x surface: *F_1,265_* = 36.2, *p* < 0.001; vision x group: *F_1,265_* = 481.2, *p* < 0.001; surface x group: *F_1,265_* = 53.8, *p* < 0.001; vision x surface x group: *F_1,265_* = 40.6, *p* < 0.001). Conversely, only having had a stroke had an effect on the CKC derived from vCoP_AP_ (*F_1,265_*= 830.7, *p* < 0.001) and the interactions vision x surface (*F_1,265_* = 33.3, *p* < 0.001), vision x group (*F_1,265_* = 8.6, *p* < 0.004), and vision x surface x group (*F_1,265_* = 14.4, *p* < 0.001). The random effects statistical outcomes are reported in Supplementary information (Table S5, S6, and S7). The collection site accounted for significant differences in CKC grand averages between cohorts and its relationship to having suffered a stroke. Post-hoc comparisons (see Figure 3) indicated that CKC strength increased with condition complexity. CKC strength was higher in stroke participants compared with controls when derived from CoP_ML_ when standing on foam, either eyes open or closed, and when derived from rCoP when standing on firm ground with eyes open.

**Figure 3.**
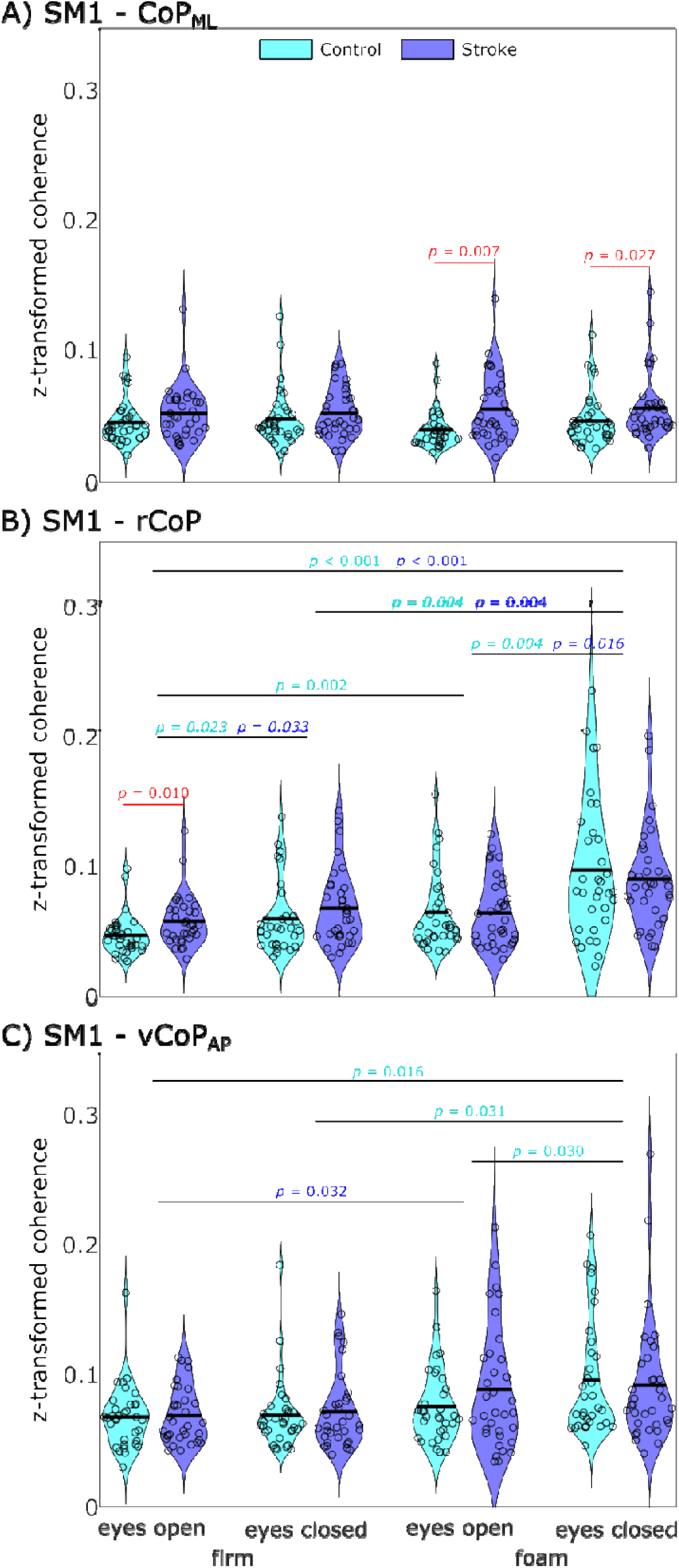
Distribution of CKC values derived from CoP_ML_ (A), rCoP (B) and vCoP_AP_ (C). Displayed are the CKC values averaged across 1–10 Hz and z-transformed (by means of the hyperbolic arctangent of the square-root). Significant differences between groups (in red) and between conditions (in cyan within the control group and in blue within the stroke group) are indicated with horizontal lines and the corresponding p-values.

### Delays

Figure 4 presents the individual values of estimated time delays for the CKC derived from the 3 CoP features (CoP_ML_, rCoP, and vCoP_AP_). The delays for |rCoP| were significantly negative, indicating that cortical activity led rCoP (control, -54 ± 65 ms, *t_23_* = -3.31, *p* < 0.001; stroke, -49 ± 84 ms, *t_25_* = -2.81, *p* = 0.005). The delays for CoP_ML_ (control, 34 ± 125 ms, *p* = 0.70; stroke, 34 ± 90 ms, *p* = 0.31) and vCoP_AP_ (control, 26 ± 89 ms, *p* = 0.24; stroke, 10 ± 122 ms, *p* = 0.79) were not significantly different from 0 for both groups.

**Figure 4.**
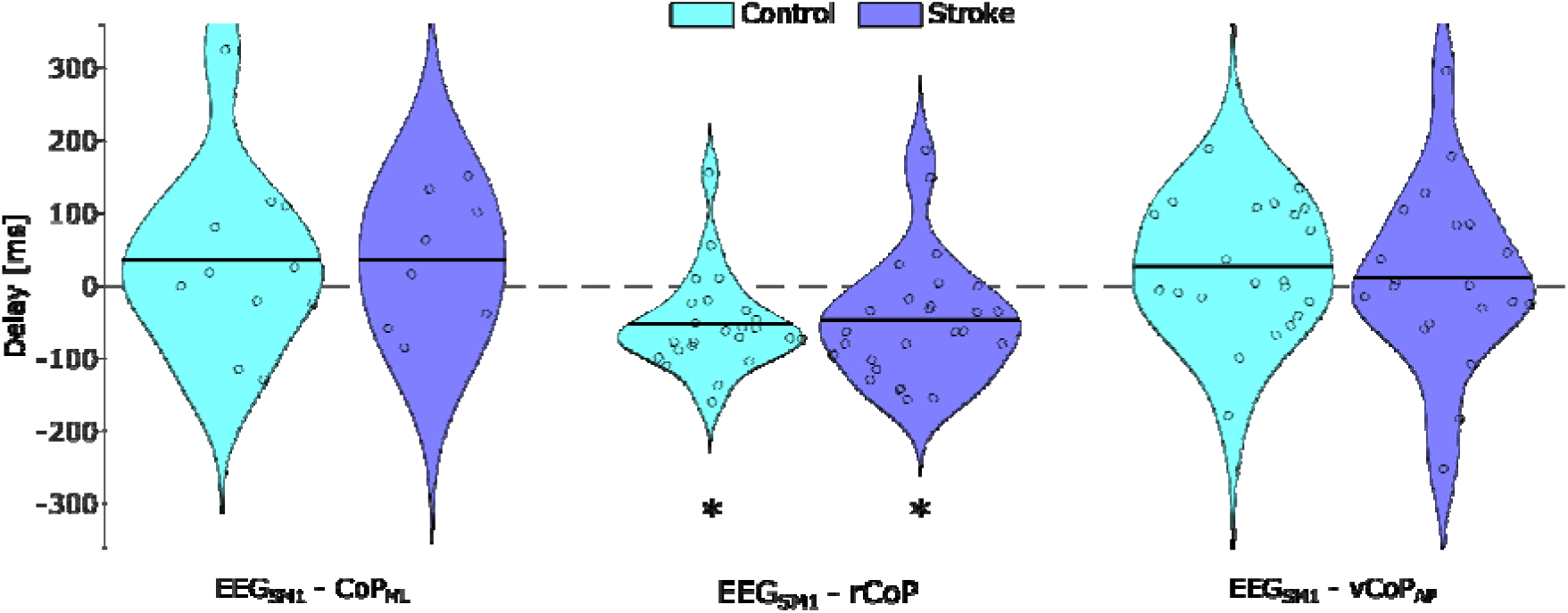
Time delay estimation underlying sway-based CKC. Positive delays indicate that the CoP feature leads cortical activity. *: delay significantly different from 0.

The delays for rCoP in the stroke group were not significantly different from controls (*z* = 0, *p* = 1).

### Behavioural relevance of CoP cortical encoding

We then sought to assess the relationship between CKC and postural sway. First, the two measures of postural sway, SD(CoP_AP_) and mean(|vCoP|), were highly correlated within both groups in all balance conditions (control, *r* = 0.86, *p* < 0.001; stroke, *r* = 0.94, *p* < 0.001). They were thus combined into a single measure of sway amplitude by averaging them after standardization. Then, behavioral relevance was assessed using the multilinear regression between the relative change in CKC strength for the 3 CoP features and sway amplitude between standing eyes open on firm ground and the other balance conditions.

Figure 5 presents the result of the multilinear regression analysis, which revealed significant associations for both groups (control, *r* = 0.48, *p* < 0.001; stroke, *r* = 0.42, *p* < 0.001). Moreover, in both cases, the weights for CKC features indicated a positive correlation between instability and CKC strength for vCoP_AP_ and rCoP. These results indicate that postural sways tend to increase when CKC derived from rCoP and vCoP_AP_ is high for healthy participants and stroke survivors. Additionally, the variance of the relative change in CKC and postural sway is represented by circles centred on the centroid of each condition with a radius equal to the average Euclidean distance between each individual and the centroid. The variance of relative change was significantly larger for stroke participants when standing on foam, either eyes open (*t* = 4.15, *p* < 0.001) or closed (*t* = 3.35, *p* = 0.001).

**Figure 5.**
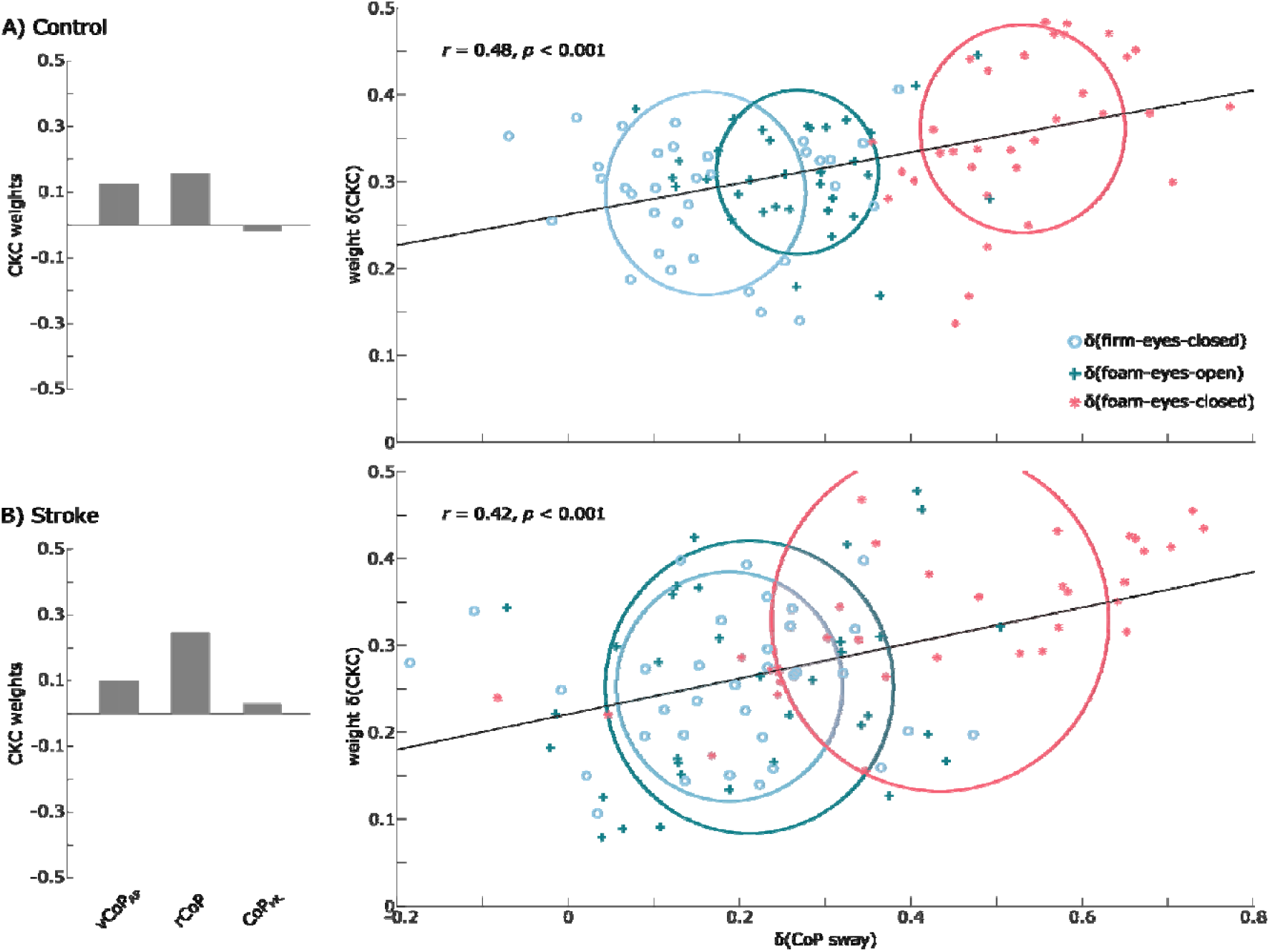
Behavioral relevance of the cortical encoding of postural sways as assessed with a multilinear regression within the control and stroke group separately. The multilinear regression was assessed between the relative change in CKC values for the 3 features and the relative change in average postural sway parameters between standing eyes open on solid ground and the other balance conditions. On the left are the weights for CKC strength identified by the multilinear regression. On the right are the scatter plots for the combination of CKC strength identified by the multilinear regression model as a function of sway amplitude. Each individual participant’s value is indicated with a circle, a cross, or a star, depending on the balance condition, and the regression line is in black. The variance in relative change for each balance condition is displayed as a circle of the corresponding color.

Given that sway-based CKC appears to be behaviorally relevant for stability, we further sought to clarify its relationship with functional mobility, as assessed by the Berg Balance Scale score. Table 3 presents the results of linear fixed-effects models for the association between CKC values and sway parameters across all balance conditions, as well as with Berg Balance Scale scores. This analysis identified a significant association when standing eyes closed on firm ground between the Berg Balance Scale score and both SD(CoP_AP_) and mean(|vCoP|). Further testing using Spearman correlation revealed a significant negative correlation for both SD(CoP_AP_) (*r* = -0.34, *p* = 0.048) and mean(|vCoP|) (*r* = -0.40, *p* = 0.020). In terms of CKC, the Berg Balance Scale scores were significantly associated with the three features (CoP_ML_, rCoP, and vCoP_AP_) when standing eyes open on firm ground and standing eyes closed on firm ground, as well as in the case of CKC derived from CoP_ML_. Spearman correlations further revealed that CKC derived from rCoP was significantly negatively correlated with the Berg Balance Scale scores (*r* = -0.37, *p* = 0.030), while CoP_ML_ and vCoP_AP_ were not significantly correlated.

**Table 3.**
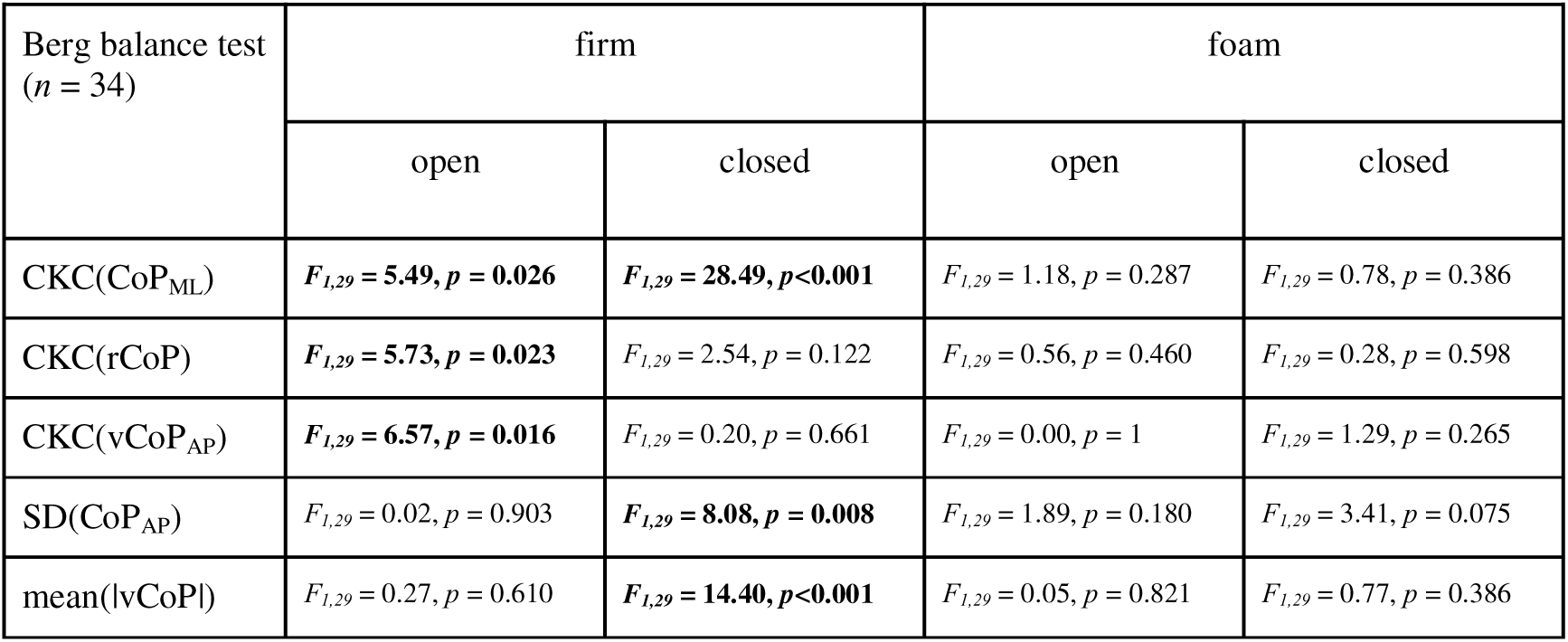
Fixed effect statistics of linear mixed effect models between CKC for each CoP feature (CoP_ML_, rCoP and vCoP_AP_), the sway parameters (SD(CoP_AP_) and mean(|vCoP|)) for each balance condition, and the Berg Balance Scale score.

## Discussion

This study aimed at characterizing the impact of a stroke on the cortical involvement in balance maintenance, evaluated with sway-based CKC. In line with previous results on a young cohort (28), sway-based CKC emerged mainly between 2 and 6 Hz, with scalp distributions suggestive of dominant sources in the lower limb area of the primary sensorimotor cortex. Here, we found significantly increased CKC during medio-lateral sway in stroke participants when standing on foam compared with healthy controls, suggesting increased cortical involvement to compensate for decreased function of the hemiparetic side.

### Impact of a stroke on balance performance

Overall, our results did not show any significant differences in postural sway (SD(CoPAP) and mean(|vCoP|)) between stroke participants and controls, except for an increase in mean(|vCoP|) when stroke participants stood on firm ground with eyes closed. However, weight-bearing asymmetry between lower limbs, a persistent impairment often exhibited by stroke survivors (11), was not quantified. This may have contributed to the absence of postural sway difference between cohorts. This may also be explained by the high functional mobility of our cohort of stroke participants, as evidenced by the cohort’s average clinical scores. Supporting this view, the clinical scores were not significantly associated with postural sway parameters, except for the Berg Balance Scale, which showed a significant association when standing with eyes closed on firm ground. Thus, this suggests that a higher Berg Balance Scale score indicates better balance performance in our cohort when standing with eyes closed, consistent with previous studies (40).

### Impact of a stroke on cortical involvement

Overall, stroke participants showed coupling between sensorimotor cortical activity and postural sway similar to that of controls across balance conditions. Similarly, no significant difference in the delay between cortical activity and postural sway was observed between cohorts, suggesting that, on average, stroke did not affect participants’ processing speed, at least with respect to the outcomes of interest.

When specifically considering coupling with medio-lateral postural sway, stroke participants showed a significant increase in CKC strength when standing on foam, with eyes open or closed, compared with controls. As no significant differences in postural sway were found between groups under these conditions, this suggests a compensatory strategy to mitigate any residual decrease in sensorimotor function on the hemiparetic side, resulting in increased cortical involvement in medio-lateral postural sway. This result is consistent with the weight-bearing asymmetry observed even in clinically well-recovered stroke survivors (11).

Interestingly, a significant increase in CKC strength was observed for CKC(rCoP) when participants stood on firm ground with their eyes open, particularly among stroke participants, suggesting greater cortical involvement in maintaining balance even under the easiest condition. Again, no significant difference in postural sway was associated with this difference in CKC strength, suggesting the presence of a compensatory strategy to maintain balance performance similar to that of healthy controls. A study by Kukkar et al. (41) did not find any difference in cortico-muscular coherence (CMC), the coupling between cortical and muscle activity, between chronic stroke participants and healthy controls during quiet standing, and even found a decrease in delta band CMC in more challenging balance conditions. This discrepancy may be explained by differences in balance performance between our cohorts: on average, participants in our study scored 53 on the Berg Balance Scale, whereas participants in their cohort scored 42, suggesting less functional recovery and compensation in their cohort. In that regard, our results are similar to what has been observed for the upper limb in stroke survivors, where CMC increased with motor recovery, and was significantly higher at a chronic stage compared to healthy controls (42).

This difference, observed in the easiest balance condition of our study, which is commonly encountered in daily life, suggests that stroke participants must continually maintain elevated levels of sensorimotor cortical activity. Several studies have suggested a link between changes in cortical activity and fatigue, a common symptom post-stroke (43–45). Although the pattern of change in cortical activity leading to fatigue remains unclear, it appears to be largely linked to increased prefrontal cortical activation (44, 45), particularly during challenging balance tasks (45). The evaluation of post-stroke fatigue and a connectivity analysis were out of the scope of the present study; thus, a link between fatigue and prefrontal cortical activation during quiet standing cannot be established. However, many tasks of daily living are performed standing; therefore, further investigation of a potential link between the two is warranted, as it could lead to balance interventions that make postural control more automatic and less tiring.

### Behavioural relevance

In terms of the behavioral relevance of CKC for balance performance, our results confirmed a positive relationship between CKC strength and the amount of postural sway. When looking at the relative increase in cortical involvement and postural sway compared to quiet standing, the correlations were moderate and significant for both the stroke and healthy participants, similar to the study by Legrand et al. (28), which found a weak positive correlation between the relative increase in postural sway and CKC strength. Overall, individuals with the greatest increase in postural sway also showed the greatest increase in cortical involvement, irrespective of group. However, stroke survivors showed significantly higher within-group variance when standing on foam, either eyes open or closed. This highlights heterogeneity in postural control within the stroke cohort, despite similar scores on the clinical tests (FMA and BBS). This greater variance could suggest the presence of several long-established compensatory mechanisms that have enabled them to achieve functional levels on par with healthy controls. Interestingly, this also shows the potential of CKC to stratify and identify stroke survivors who might still benefit from further balance rehabilitation. This would first require defining what constitutes normative CKC levels in a large cohort of healthy controls, thus allowing a direct comparison for CKC survivors and constituting a potential rehabilitation target.

Similarly, CKC derived from rCoP was significantly associated with the Berg Balance scores when standing on firm ground, eyes open. This suggests that the scores, in part, reflect the level of cortical involvement required to maintain balance.

### Limitations

No attempt was made to localize the cortical sources of CKC to account for the heterogeneity of lesions or compensations of our stroke cohort. In question is the accuracy of source localization with EEG, which is at best D∼D1 cm for focal cortical sources (46). Moreover, the contributions of close-by areas cannot be resolved owing to the ill-posed nature of the inverse solution.

Not all participants in either group exhibited significant CKC. And even when CKC was detected, its amplitude was well below the theoretical maximum of 1. However, this was expected, as the cerebral cortex is thought to be involved in postural control only intermittently. Overall, the proportion of participants with significant CKC was comparable to that reported in a previous study (28).

The phase delays underlying CKC exhibited substantial inter-individual variability, with some individual values showing the opposite sign to the group average. This variation is most likely attributable to the low amplitude of CKC, which leads to inaccuracies in the estimation of individual values. Alternatively, we cannot exclude the possibility that there is inter-individual variability in the way the brain monitors and controls CoP fluctuations.

The Berg Balance and Fugl-Meyer tests suffer from a ceiling effect that may have concealed differences in ability between highly performing stroke survivors (47, 48).

Significant variability was found between cohorts, which affected the impact of the balance conditions and of having suffered a stroke on postural sway and CKC. This was expected, as the results are based on a small sample of stroke survivors, mainly at a chronic stage, from three distinct cohorts, collected with different equipment, thus limiting their generalizability. Still, significant main effects from the balance condition and of having suffered a stroke were still apparent. Further research should explore the evolution of CKC and its correlation with functional aspects at the acute and subacute phases of stroke.

## Conclusion

Despite the functional consequences of a stroke on postural control, which in turn compromises autonomy and increases the risk of falls in stroke survivors (3–5), the majority of current studies on cortical alterations in stroke patients have concentrated on upper limb assessments, neglecting comprehensive investigations into postural control. Our results show that, even in highly functional stroke survivors, as measured by clinical scores, differences in cortical involvement persist relative to healthy controls. The increased use of cortical resources in chronic stroke indicates a shift from more autonomous, or spinally-mediated, postural control to a more cortically mediated form of motor control. While this may represent a compensatory strategy to achieve performance comparable to healthy controls, it may also result in greater fatigue during daily life (45). The level of cortical involvement was only weakly correlated with clinical scores, suggesting that CKC could be used as a biomarker to track progress beyond traditional functional recovery as measured by clinical scores. Thus, balance training might be beneficial, especially as there is now a robust evidence base for stroke rehabilitation even at a chronic stage (49, 50).

## Acknowledgements

This publication has emanated from research supported under the European Union’s Horizon 2020 research and innovation programme under the Marie Skłodowska-Curie grant agreement No 101034252.

## Data availability

The data that support the findings of this study are available from the corresponding author upon reasonable request.

