## Supporting Information for "Cortical involvement in stroke survivors for balance maintenance"

**This PDF file includes:**

Table S1 to S7

*Table S1. Stroke participants demographics and stroke details.*

| # | Site | Age | Sex | Type of stroke | Type of lesion |
| --- | --- | --- | --- | --- | --- |
| 01 | Ferrara | 64 | M | haemorrhagic | Impression on the right lateral ventricle and the third ventricle |
| 02 | Ferrara | 78 | M | ischemic | Right subcortical frontal region. |
| 03 | Ferrara | 59 | M | ischemic | Cortical areas in the right hemisphere. |
| 04 | Ferrara | 76 | M | ischemic | Hypodensity of the posterior limb of the right internal capsule. |
| 05 | Ferrara | 51 | M | haemorrhagic | Left temporal insular corona radiata and centrum semiovale |
| 06 | Ferrara | 72 | M | ischemic | Left bulbar lesion |
| 07 | Ferrara | 55 | F | ischemic | Left capsulo-lenticular region extending to the corona radiata. |
| 08 | Ferrara | 65 | F | ischemic | Right fronto-temporo-insular lesion. |
| 09 | Ferrara | 65 | M | haemorrhagic | Right capsulo-thalamic hematoma |
| 10 | Ferrara | 54 | M | haemorrhagic | Right insular and capsulo-lenticular region. |
| 11 | Ferrara | 61 | M | ischemic | Left lenticulo-capsular region. |
| 12 | Ferrara | 40 | M | haemorrhagic | Right intraparenchymal |
| 13 | Ferrara | 57 | F | ischemic | Right caudate-capsulo-lenticular region |
| 14 | Ferrara | 52 | M | ischemic | Right fronto-insular region |
| 15 | UCD | 35 | F | haemorrhagic | Double sub arachnoid |
| 16 | UCD | 40 | F | ischemic | Brain tumor on the right hemisphere |
| 17 | UCD | 45 | F | ischemic | Left hemisphere |
| 18 | UCD | 50 | F | ischemic | Right middle cerebellum |
| 19 | UCD | 55 | M | ischemic | Right hemisphere |
| 20 | UCD | 51 | M | ischemic | right middle cerebellar artery |
| 21 | UCD | 71 | M | ischemic | Right hemisphere |
| 22 | UCD | 40 | F | ischemic | clot on left parietal and frontal lobes |
| 23 | Erasme | 73 | M | ischemic | Left temporal and frontal lobes |
| 24 | Erasme | 54 | M | ischemic | Left pons |
| 25 | Erasme | 60 | F | ischemic | Right junctional territory |
| 26 | Erasme | 87 | M | ischemic | Left middle cerebral artery |
| 27 | Erasme | 35 | M | haemorrhagic | Right parietal lobe |
| 28 | Erasme | 93 | F | ischemic | Anterior inferior cerebellar artery |
| 29 | Erasme | 88 | F | ischemic | Left parietal and occipital lobes |
| 30 | Erasme | 59 | M | ischemic | Right middle cerebral artery |
| 31 | Erasme | 57 | M | haemorrhagic | Right capsulo-lenticular |
| 32 | Erasme | 65 | M | ischemic | Vertebrobasilar arteries |
| 33 | Erasme | 73 | M | haemorrhagic | Right capsulo-lenticular |
| 34 | Erasme | 60 | M | ischemic | Right frontal, postcentral and occipital lobes |

*Table S2. Random effects statistics for SD(CoP_ML_)*

|  |  | Estimate | SE | DF | *p* |
| --- | --- | --- | --- | --- | --- |
| Ferrara | Intercept | 0.62 | 0.27 | 5.95 | 0.060 |
| Ferrara | vision_open | -0.16 | 0.07 | 1.37 | 0.201 |
| Ferrara | surface_solid | -0.27 | 0.12 | 3.28 | 0.096 |
| Ferrara | group_stroke | -0.21 | 0.09 | 0.62 | 0.366 |
| Ferrara | vision_open:surface_solid | 0.10 | 0.04 | 0.50 | 0.416 |
| Ferrara | vision_open:group_stroke | 0.01 | 0.00 | 0.00 | 0.986 |
| Ferrara | surface_solid:group_stroke | -0.05 | 0.02 | 0.13 | 0.727 |
| Ferrara | vision_open:surface_solid:group_stroke | -0.01 | 0.01 | 0.01 | 0.960 |
| UCD | Intercept | 1.98 | 0.17 | 126.08 | **<0.001** |
| UCD | vision_open | -0.52 | 0.04 | 1.52 | **0.017** |
| UCD | surface_solid | -0.85 | 0.07 | 3.93 | **<0.001** |
| UCD | group_stroke | -0.66 | 0.05 | 1.35 | **0.025** |
| UCD | vision_open:surface_solid | 0.30 | 0.03 | 0.53 | 0.170 |
| UCD | vision_open:group_stroke | 0.02 | 0.00 | 0.00 | 0.981 |
| UCD | surface_solid:group_stroke | -0.16 | 0.01 | 0.11 | 0.625 |
| UCD | vision_open:surface_solid:group_stroke | -0.04 | 0.00 | 0.01 | 0.943 |
| Erasme | Intercept | 3.06 | 0.11 | 147.94 | **<0.001** |
| Erasme | vision_open | -0.80 | 0.03 | 1.45 | **0.006** |
| Erasme | surface_solid | -1.31 | 0.05 | 3.55 | **<0.001** |
| Erasme | group_stroke | -1.02 | 0.04 | 1.41 | **0.007** |
| Erasme | vision_open:surface_solid | 0.47 | 0.02 | 0.52 | 0.114 |
| Erasme | vision_open:group_stroke | 0.04 | 0.00 | 0.00 | 0.979 |
| Erasme | surface_solid:group_stroke | -0.24 | 0.01 | 0.11 | 0.569 |
| Erasme | vision_open:surface_solid:group_stroke | -0.07 | 0.00 | 0.01 | 0.935 |

*Table S3. Random effects statistics for SD(CoP_AP_)*

|  |  | Estimate | SE | DF | *p* |
| --- | --- | --- | --- | --- | --- |
| Ferrara | Intercept | 2.59 | 0.23 | 12.49 | **<0.001** |
| Ferrara | vision_open | -1.48 | 0.13 | 4.61 | **<0.001** |
| Ferrara | surface_solid | -1.21 | 0.11 | 3.54 | **<0.001** |
| Ferrara | group_stroke | -0.09 | 0.01 | 0.01 | 0.936 |
| Ferrara | vision_open:surface_solid | 1.00 | 0.09 | 2.33 | **0.005** |
| Ferrara | vision_open:group_stroke | 0.12 | 0.01 | 0.03 | 0.874 |
| Ferrara | surface_solid:group_stroke | -0.64 | 0.06 | 1.05 | 0.051 |
| Ferrara | vision_open:surface_solid:group_stroke | 0.08 | 0.01 | 0.01 | 0.927 |
| UCD | Intercept | 3.92 | 0.20 | 74.8 | **<0.001** |
| UCD | vision_open | -2.23 | 0.12 | 5.77 | **<0.001** |
| UCD | surface_solid | -1.84 | 0.09 | 3.54 | **<0.001** |
| UCD | group_stroke | -0.14 | 0.01 | 0.01 | 0.926 |
| UCD | vision_open:surface_solid | 1.51 | 0.08 | 2.52 | **<0.001** |
| UCD | vision_open:group_stroke | 0.18 | 0.01 | 0.03 | 0.855 |
| UCD | surface_solid:group_stroke | -0.97 | 0.05 | 0.93 | **0.040** |
| UCD | vision_open:surface_solid:group_stroke | 0.12 | 0.01 | 0.01 | 0.921 |
| Erasme | Intercept | 5.16 | 0.15 | 161.39 | **<0.001** |
| Erasme | vision_open | -2.94 | 0.08 | 4.65 | **<0.001** |
| Erasme | surface_solid | -2.42 | 0.07 | 3.18 | **<0.001** |
| Erasme | group_stroke | -0.18 | 0.01 | 0.01 | 0.918 |
| Erasme | vision_open:surface_solid | 1.99 | 0.06 | 2.29 | **<0.001** |
| Erasme | vision_open:group_stroke | 0.23 | 0.01 | 0.03 | 0.842 |
| Erasme | surface_solid:group_stroke | -1.28 | 0.04 | 0.91 | **0.024** |
| Erasme | vision_open:surface_solid:group_stroke | 0.16 | 0.00 | 0.01 | 0.913 |

*Table S4. Random effects statistics for mean(|vCoP|)*

|  |  | Estimate | SE | DF | *p* |
| --- | --- | --- | --- | --- | --- |
| Ferrara | Intercept | 14.52 | 1.52 | 9.79 | **<0.001** |
| Ferrara | vision_open | -8.09 | 0.85 | 3.24 | **0.002** |
| Ferrara | surface_solid | -6.79 | 0.80 | 0.65 | 0.155 |
| Ferrara | group_stroke | 1.78 | 0.32 | 0.04 | 0.841 |
| Ferrara | vision_open:surface_solid | 5.46 | 0.58 | 1.24 | **0.042** |
| Ferrara | vision_open:group_stroke | -0.03 | 0.07 | 0.00 | 0.991 |
| Ferrara | surface_solid:group_stroke | -2.94 | 0.46 | 0.09 | 0.714 |
| Ferrara | vision_open:surface_solid:group_stroke | 0.08 | 0.06 | 0.00 | 0.991 |
| UCD | Intercept | 19.97 | 1.43 | 43.99 | **<0.001** |
| UCD | vision_open | -11.07 | 0.80 | 3.87 | **<0.001** |
| UCD | surface_solid | -9.67 | 0.74 | 0.75 | 0.092 |
| UCD | group_stroke | 2.18 | 0.33 | 0.04 | 0.855 |
| UCD | vision_open:surface_solid | 7.58 | 0.54 | 1.50 | **0.014** |
| UCD | vision_open:group_stroke | 0.03 | 0.07 | 0.00 | 0.992 |
| UCD | surface_solid:group_stroke | -4.36 | 0.44 | 0.08 | 0.707 |
| UCD | vision_open:surface_solid:group_stroke | 0.17 | 0.06 | 0.00 | 0.989 |
| Erasme | Intercept | 27.71 | 1.07 | 71.85 | **<0.001** |
| Erasme | vision_open | -15.41 | 0.60 | 2.69 | **<0.001** |
| Erasme | surface_solid | -13.10 | 0.59 | 0.36 | 0.224 |
| Erasme | group_stroke | 3.27 | 0.31 | 0.03 | 0.872 |
| Erasme | vision_open:surface_solid | 10.45 | 0.41 | 1.27 | **0.011** |
| Erasme | vision_open:group_stroke | -0.03 | 0.07 | 0.00 | 0.992 |
| Erasme | surface_solid:group_stroke | -5.75 | 0.38 | 0.06 | 0.766 |
| Erasme | vision_open:surface_solid:group_stroke | 0.18 | 0.06 | 0.00 | 0.989 |

*Table S5. Random effects statistics for CKC(rCoP)*

|  |  | Estimate | SE | DF | *p* |
| --- | --- | --- | --- | --- | --- |
| Ferrara | Intercept | 7.54 | 7.72 | 1.87 | 0.437 |
| Ferrara | vision_open | 0.63 | 0.64 | 0.10 | 0.831 |
| Ferrara | surface_solid | 1.11 | 1.13 | 0.31 | 0.683 |
| Ferrara | group_stroke | -7.33 | 7.49 | 1.74 | 0.444 |
| Ferrara | vision_open:surface_solid | -1.26 | 1.29 | 0.36 | 0.659 |
| Ferrara | vision_open:group_stroke | -0.62 | 0.64 | 0.10 | 0.832 |
| Ferrara | surface_solid:group_stroke | -0.65 | 0.66 | 0.10 | 0.830 |
| Ferrara | vision_open:surface_solid:group_stroke | 1.15 | 1.18 | 0.30 | 0.688 |
| UCD | Intercept | -13.66 | 0.37 | 176.95 | **<0.001** |
| UCD | vision_open | -1.13 | 0.03 | 0.11 | 0.552 |
| UCD | surface_solid | -2.00 | 0.05 | 0.34 | 0.197 |
| UCD | group_stroke | 13.27 | 0.36 | 194.10 | **<0.001** |
| UCD | vision_open:surface_solid | 2.29 | 0.06 | 0.45 | 0.127 |
| UCD | vision_open:group_stroke | 1.13 | 0.03 | 0.11 | 0.557 |
| UCD | surface_solid:group_stroke | 1.18 | 0.03 | 0.12 | 0.534 |
| UCD | vision_open:surface_solid:group_stroke | -2.09 | 0.06 | 0.38 | 0.172 |
| Erasme | Intercept | -15.74 | 0.25 | 165.22 | **<0.001** |
| Erasme | vision_open | -1.31 | 0.02 | 0.11 | 0.523 |
| Erasme | surface_solid | -2.31 | 0.04 | 0.34 | 0.168 |
| Erasme | group_stroke | 15.29 | 0.24 | 241.30 | **<0.001** |
| Erasme | vision_open:surface_solid | 2.64 | 0.04 | 0.44 | 0.105 |
| Erasme | vision_open:group_stroke | 1.30 | 0.02 | 0.11 | 0.527 |
| Erasme | surface_solid:group_stroke | 1.36 | 0.02 | 0.12 | 0.504 |
| Erasme | vision_open:surface_solid:group_stroke | -2.41 | 0.04 | 0.37 | 0.147 |

*Table S6. Random effects statistics for CKC(CoP_ML_)*

|  |  | Estimate | SE | DF | *p* |
| --- | --- | --- | --- | --- | --- |
| Ferrara | Intercept | 7.11 | 7.35 | 1.60 | 0.457 |
| Ferrara | vision_open | -2.04 | 2.11 | 0.47 | 0.618 |
| Ferrara | surface_solid | 0.80 | 0.84 | 0.12 | 0.817 |
| Ferrara | group_stroke | -6.62 | 6.84 | 1.36 | 0.474 |
| Ferrara | vision_open:surface_solid | 0.51 | 0.53 | 0.05 | 0.903 |
| Ferrara | vision_open:group_stroke | 2.05 | 2.12 | 0.48 | 0.617 |
| Ferrara | surface_solid:group_stroke | -0.56 | 0.59 | 0.06 | 0.888 |
| Ferrara | vision_open:surface_solid:group_stroke | -0.56 | 0.58 | 0.06 | 0.889 |
| UCD | Intercept | -16.29 | 0.39 | 52.99 | **<0.001** |
| UCD | vision_open | 4.65 | 0.11 | 0.93 | **0.019** |
| UCD | surface_solid | -1.73 | 0.15 | 0.15 | 0.543 |
| UCD | group_stroke | 15.09 | 0.37 | 12.30 | **<0.001** |
| UCD | vision_open:surface_solid | -1.19 | 0.04 | 0.02 | 0.872 |
| UCD | vision_open:group_stroke | -4.68 | 0.11 | 0.92 | **0.020** |
| UCD | surface_solid:group_stroke | 1.36 | 0.13 | 0.12 | 0.625 |
| UCD | vision_open:surface_solid:group_stroke | 1.24 | 0.05 | 0.04 | 0.791 |
| Erasme | Intercept | -18.04 | 0.27 | 59.54 | **<0.001** |
| Erasme | vision_open | 5.20 | 0.08 | 0.56 | 0.062 |
| Erasme | surface_solid | -2.13 | 0.14 | 0.14 | 0.543 |
| Erasme | group_stroke | 16.89 | 0.27 | 3.11 | **<0.001** |
| Erasme | vision_open:surface_solid | -1.28 | 0.04 | 0.02 | 0.897 |
| Erasme | vision_open:group_stroke | -5.22 | 0.08 | 0.77 | **0.025** |
| Erasme | surface_solid:group_stroke | 1.33 | 0.13 | 0.11 | 0.626 |
| Erasme | vision_open:surface_solid:group_stroke | 1.45 | 0.05 | 0.03 | 0.835 |

*Table S7. Random effects statistics for CKC(vCoP_AP_)*

|  |  | Estimate | SE | DF | *p* |
| --- | --- | --- | --- | --- | --- |
| Ferrara | Intercept | 7.47 | 7.50 | 1.57 | 0.448 |
| Ferrara | vision_open | 0.24 | 0.25 | 0.00 | 0.992 |
| Ferrara | surface_solid | 0.53 | 0.52 | 0.01 | 0.975 |
| Ferrara | group_stroke | -7.69 | 7.57 | 1.62 | 0.437 |
| Ferrara | vision_open:surface_solid | 0.70 | 0.69 | 0.01 | 0.967 |
| Ferrara | vision_open:group_stroke | -0.55 | 0.57 | 0.01 | 0.976 |
| Ferrara | surface_solid:group_stroke | 0.02 | 0.20 | 0.03 | 0.983 |
| Ferrara | vision_open:surface_solid:group_stroke | -0.26 | 0.26 | 0.00 | 0.994 |
| UCD | Intercept | -16.69 | 0.51 | 56.14 | **<0.001** |
| UCD | vision_open | -0.35 | 0.14 | 0.18 | 0.650 |
| UCD | surface_solid | -0.81 | 0.25 | 0.58 | 0.317 |
| UCD | group_stroke | 16.36 | 0.65 | 4.10 | **<0.001** |
| UCD | vision_open:surface_solid | -1.49 | 0.06 | 0.03 | 0.854 |
| UCD | vision_open:group_stroke | 1.31 | 0.08 | 0.05 | 0.791 |
| UCD | surface_solid:group_stroke | 0.24 | 0.21 | 0.39 | 0.606 |
| UCD | vision_open:surface_solid:group_stroke | 0.53 | 0.04 | 0.02 | 0.912 |
| Erasme | Intercept | -16.68 | 0.35 | 40.90 | **<0.001** |
| Erasme | vision_open | -0.70 | 0.13 | 0.18 | 0.570 |
| Erasme | surface_solid | -1.47 | 0.25 | 0.57 | 0.225 |
| Erasme | group_stroke | 17.80 | 0.56 | 2.15 | **0.001** |
| Erasme | vision_open:surface_solid | -1.62 | 0.05 | 0.03 | 0.841 |
| Erasme | vision_open:group_stroke | 1.16 | 0.07 | 0.05 | 0.779 |
| Erasme | surface_solid:group_stroke | -0.28 | 0.20 | 0.40 | 0.570 |
| Erasme | vision_open:surface_solid:group_stroke | 0.64 | 0.04 | 0.02 | 0.908 |
